# Differential benefits for temporal intervals and line segments with feedback and relevance to new learning with transfer effects

**DOI:** 10.64898/2026.07.26.740865

**Authors:** Jiaxuan Teng, Arne D. Ekstrom, Eve A. Isham

**Affiliations:** Department of Psychology, University of Arizona

**Keywords:** temporal learning, spatial learning, magnitude learning, error correction, interval timing, feedback

## Abstract

Learning to accurately estimate temporal intervals and spatial magnitudes is important to everyday behavior, yet few studies have directly compared how learning unfolds across these domains. Theoretical models of temporal interval estimation provide conflicting accounts of how this learning process might occur and how it might differ from other entities like spatial line estimation. In the present study, participants completed temporal interval and line-length production tasks under feedback or no-feedback conditions during a training phase. They were then tested without feedback and subsequently presented with untrained temporal intervals and line lengths to examine whether learning generalized to novel magnitudes. We found that both temporal and spatial production exhibited similar learning trajectories, characterized by rapid initial improvements, followed by asymptotic performance. Despite similar learning trajectories, however, spatial production reached higher levels of accuracy and precision faster and in a more sustained manner than temporal production. Feedback facilitated learning in both domains, with improvements in precision emerging early during training and improvements in accuracy becoming evident during the subsequent test phase. These benefits persisted into the transfer phase, particularly for spatial production, indicating that feedback-supported learning generalized to untrained magnitudes. Transfer performance also reflected systematic biases in magnitude estimation, consistent with a central tendency effect, with shorter magnitudes tending to be overproduced and longer magnitudes underestimated. Together, these findings demonstrate that temporal and spatial magnitude learning share some common and some dissociable learning dynamics, providing new insight into the shared and distinct mechanisms underlying magnitude learning and highlighting the important role of feedback in promoting learning and transfer.

## Introduction

Unlike other sensory modalities such as vision or hearing, time perception lacks a dedicated sensory system. Instead, it arises from complex cognitive and neural processes that integrates information beyond direct physical input [1, 2]. Yet, timing and time perception are critical for survival, although the extent to which we can encode, represent, and maintain temporal information, as well as monitor and correct temporal errors, remains unclear. These differences raise important questions about how individuals achieve proficiency such that they develop accurate (how close one is to the target interval) and precise timing (how consistent performance is across trials). One goal of our proposed study, therefore, is to determine how feedback benefits temporal interval learning.

The literature on temporal awareness suggest that humans can monitor their timing errors and evaluate performance even in the absence of external feedback [3–7]. For example, Akdoğan and Balcı [3] found that participants were metacognitively aware of both the direction and magnitude of their timing errors. Specifically, participants reported lower confidence when errors were larger and were able to distinguish whether their responses were too short or too long. Although this internal monitoring ability may provide some benefits for temporal accuracy, it is often insufficient for effective and stable error correction. Prior work has shown that trial-by-trial feedback improves accuracy and reduces systematic biases in temporal tasks [8, 9]. Despite these benefits, it remains unclear how feedback shapes learning over time. In particular, it is unknown whether improvements occur rapidly early in training and/or continue gradually with extended exposure.

There are several models of temporal interval learning that provide different perspectives on how feedback benefits learning. One line of research suggests that temporal learning can occur rapidly, sometimes even after a single trial [10, 11]. Simen et al. [10] proposed a neural integration model that includes a rapid duration-learning mechanism, showing that individuals can calibrate their timing from a single trial. Consistent with this idea, Çavdaroğlu et al. [11] further demonstrated that optimal timing behavior could be acquired quickly when reinforced by reward, with no additional improvement or worsening observed over multiple sessions. Together, these models support the perspective that learning of temporal information is rapid.

Alternative to the rapid single-trial learning, the Learning-to-Time (LeT) theory [12–14], which conceptualizes temporal learning as a form of reinforcement learning, predicts a slower and more gradual learning process. According to LeT, organisms produce accurate timed responses by forming associations between a sequence of internal states that unfold during a timed interval and the delivery of reinforcement. These internal states must be differentiated over time through repeated exposure and feedback (i.e., statistical learning); therefore, the learning process requires many trials for accurate state-reinforcement associations to emerge. The gradual formation of associations between internal states and reinforcement predicts a slow and steady trajectory of improvement, suggesting that extended training is necessary for learning to develop into proficient and accurate timing. Together, these perspectives raise a critical question: does feedback-driven temporal learning reflect rapid early adjustments, or does it continue to improve gradually with more training?

In addition to learning specific trained intervals, an important aspect of temporal learning is the ability to generalize to novel intervals. Reinforcement-based accounts such as Learning-to-Time (LeT) propose that timing behavior is shaped through associative learning tied to specific trained durations, potentially leading to limited transfer to untrained intervals. In contrast, Scalar Expectancy Theory (SET) posits a more general timing mechanism that operates across durations and supports scalar properties of timing behavior [15, 16]. Thus, examining transfer to untrained intervals provides insight into how temporal learning generalizes beyond the trained duration.

In contrast to time, spatial learning benefits from visual information, which may make feedback more effective in the spatial than in the temporal domain. Prominent theoretical models offer competing perspectives on whether these differences reflect distinct underlying cognitive mechanisms for processing space and time. Some theoretical perspectives propose that time and space share common processing mechanisms. For example, the Theory of Magnitude (ATOM [17]) suggests a unified system for representing magnitude information, including space, time, and number. Similarly, the Mental Time Line (MTL) theory posits that temporal information is spatially represented, with shorter durations mapped to the left and longer durations to the right [18–20]. Yet, other theoretical models suggest that aspects of space and time may, in some situations, dissociate from each other [21–31]. For example, evidence from decision-making research suggests that temporal and spatial discounting appear largely independent at the individual level, such that individuals who strongly discounted future temporal rewards did not necessarily strongly discount spatial rewards and vice versa [26]. Therefore, another goal of the present study was to compare the effects of feedback on learning and performance in the spatial domain, in part to test how space and time learning differ and also to provide a comparison test case for time.

In a recent study [32], participants completed a brief training session followed by assessments of learning specificity and transfer across temporal and spatial magnitudes. Although that study primarily examined metacognitive awareness of temporal errors, findings related to feedback revealed that temporal performance remained less accurate and more variable than spatial performance, despite participants demonstrating awareness of their errors. Given the brevity of the training phase (i.e., 4 trials), it remains unclear whether these effects reflect stable learning mechanisms or are driven by early-stage adjustments that emerge during initial exposure.

The present study extends this work by examining how temporal and spatial performance evolve over an extended period of training, allowing us to better understand the durable effects of feedback. Rather than focusing solely on end-point differences, we characterized learning trajectories to determine how accuracy and precision change across repeated trials and whether improvements continue, stabilize, or plateau over time. Participants completed temporal and line-length production tasks under conditions with and without explicit feedback and were subsequently tested on both trained and untrained magnitudes. This design allowed us to examine not only the effects of feedback on learning but also the extent to which learning generalized beyond the trained interval or distance.

We propose to test three primary hypotheses: 1) explicit feedback will facilitate improvements in both accuracy and precision relative to the no feedback condition 2) if temporal learning follows reinforcement-based accounts such as Learning-to-Time (LeT), improvements should emerge gradually across training. In contrast, if temporal learning reflects rapid adaptation processes [10, 11], performance should show substantial early gains followed by later stabilization. 3) spatial performance will be more accurate and precise than temporal performance due to the presence of visual information and feedback.

## Methods

### Participants

We tested sixty-one participants (40 females, 19 males, and 2 others; mean age = 19.3). Participants received class credit for their participation. Based on the exclusion criteria described in the Data Analysis section, six participants were excluded, resulting in a final sample of 55 participants (37 females, 17 males, and 1 other; mean age = 19.3). The study protocol was approved by the University of Arizona Institutional Review Board. Participants were recruited between September 2024 and February 2025. All participants provided written informed consent prior to participation.

### Procedure

Participants completed both the time interval and line length production tasks in separate blocks, with the order of the blocks counterbalanced across participants. The tasks were programmed in jsPsych [33]. In the time task, participants clicked the mouse twice to indicate the onset and offset of a target duration of 2.8 seconds (adapted from [32, 34]. Each click produced a corresponding “X” on the screen to confirm the response (see [32]), with placement determined by the participant’s mouse position. The produced duration was defined as the time interval between the two clicks. In the line-length production task, participants similarly clicked the mouse twice to indicate the beginning and end of a 2.8-centimeter line. Each click generated a corresponding “X” on the screen, and the distance between the two X’s was recorded as the produced line length.

The experimental procedure and trial structure are depicted in Fig 1. Participants were randomly assigned to either the no feedback condition or explicit feedback condition. Prior to each task, participants completed 10 practice trials without feedback to familiarize themselves with the modality, producing one-second intervals in the time task and one-centimeter lines in the line length task.

**Fig 1.**
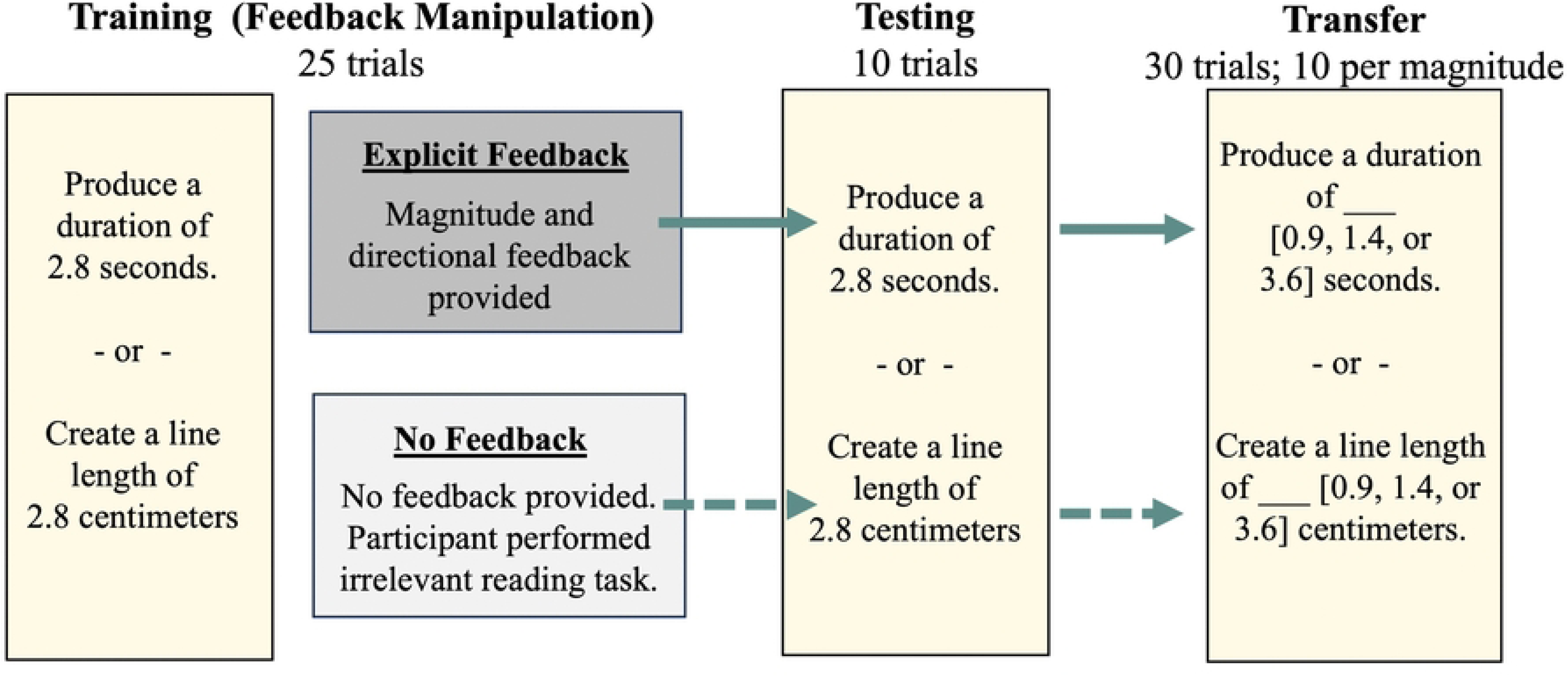
Experimental design and task flow. During the training phase, participants performed either a time production or line-length production task by pressing the computer mouse twice to mark the beginning and end of a temporal interval or line length. Participants in the explicit feedback condition received performance feedback from the computer, while those in the no feedback condition were not informed of their performance errors and instead completed a brief, unrelated reading task after each trial. In the testing phase, participants produced the same magnitude (2.8 units) without any feedback. In the final transfer phase, participants produced untrained magnitudes (0.9, 1.4, or 3.6 units). A mandatory five-minute break was provided between each phase.

#### Training Phase

During the training phase, participants completed 25 trials of either the time production task or the line-length production task. In both tasks, the produced durations and lengths were recorded by the computer, and the errors were computed for each trial. Participants in the explicit feedback condition received written feedback from the computer regarding the direction and magnitude errors (e.g., “Your judgment was 120 milliseconds shorter than the actual duration.”). On the other hand, participants in the no feedback condition were instructed to read irrelevant quotes before preceding to the next trial.

#### Testing Phase

After a brief 5-minute break, participants completed 10 trials of the production task using the same target magnitude (2.8 units) to assess the effects of training. No explicit feedback was provided in either condition. As in the training phase, participants clicked the mouse twice to indicate the onset and offset of the target duration (time task) or the start and end points of the target length (line-length task).

#### Transfer Phase

To evaluate the generalization of learning, participants then completed a transfer phase, during which they performed a production task using three untrained novel magnitudes (0.9, 1.4, and 3.6 units). These values were selected to span a range of intervals relative to the trained target (2.8 units), including both subsecond and suprasecond durations. This phase allowed us to assess whether learning generalized beyond the trained interval and whether performance varied across magnitudes.

### Data analysis

The exclusion criteria were as follows: participants who failed to follow task instructions were excluded from the analyses. In addition, data outside of 2.5 standard deviations of the group mean were considered outliers and excluded from the analyses. Accuracy was assessed using the average ratio across trials in the testing and transfer phases. The ratio was calculated as the quotient of the produced duration to the target duration. A ratio of 1 indicates that the produced duration is the same as the instructed target duration. Ratios greater than 1 reflect overproduction, while ratios less than 1 indicate underproduction, thus capturing the direction of error. Consistency of performance (precision) was quantified using the coefficient of variation (CV), computed as the standard deviation divided by the mean of the produced durations. Higher CV values indicate greater variability and lower precision. During the training phase, the 25 trials were further divided into five trial bins to examine changes in performance and learning trajectories over time.

Separate analyses were conducted for the Training, Testing, and Transfer Phases. For the training Phase, learning trajectories across trial bins were analyzed using linear mixed-effects models (LMMs) with trial bin, modality (time vs. line length), and feedback condition (explicit feedback vs. no feedback) as fixed effects and participant as a random effect. Separate models were conducted for ratio and CV to examine how performance changed across training and whether learning trajectories differed as a function of modality and feedback condition. Additional curve-fitting analyses compared logarithmic, linear, and quadratic functions to characterize patterns of learning across training and allow us to test rates of learning with and without feedback. For the testing phase, separate LMMs examined the effects of modality and feedback condition on ratio and CV measures. For the transfer phase, separate LMMs were conducted for ratio and CV, with transfer magnitude (0.9, 1.4, and 3.6) and modality included as within-subject factors and feedback condition included as a between-subjects factor. Participant was included as a random intercept to account for repeated observations within individuals.

### Open science practices statement

#### Data availability

The data and materials for all experiments are available at https://cat.lab.arizona.edu/.

## Results

### Feedback differentially affects improvements in temporal and spatial learning accuracy

To examine learning trajectories during the training block, the 25 training trials were divided into five trial bins, with each bin containing five trials. A linear mixed-effects model (LMM) was conducted to examine whether performance changed across trial bins and whether learning differed as a function of modality (time vs. space) or feedback (present or not) condition (Fig 2A). A follow-up model-fitting analysis was subsequently conducted to further characterize learning trajectories across training. We focused first on the temporal and spatial ratios as measures of accuracy.

**Fig 2.**
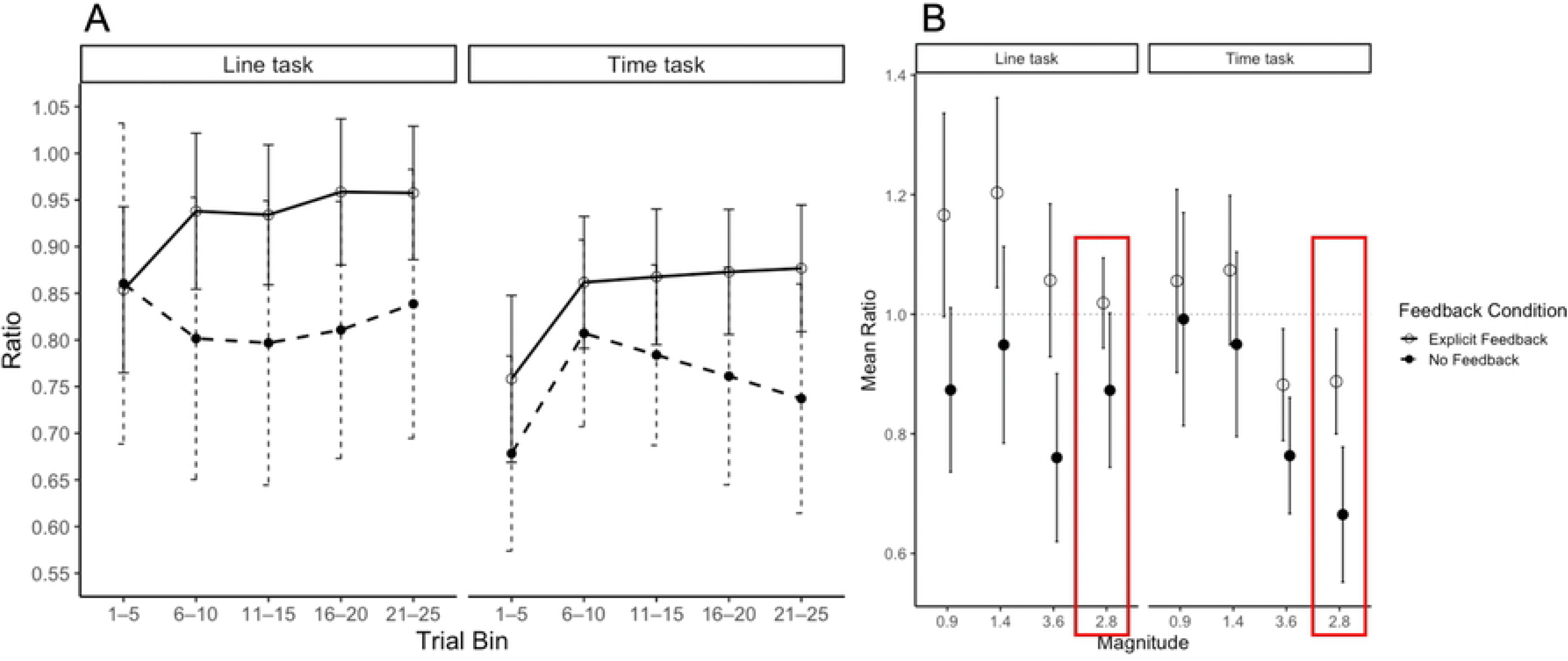
Accuracy (ratio) during learning and transfer. (A) Mean ratio across training trial bins as a function of modality and feedback condition. The line task is shown on the left and the time task on the right. (B) Mean ratio during the transfer phase across magnitudes as a function of modality and feedback condition. Performance at the trained magnitude (2.8) is highlighted by the red box and displayed alongside the untrained transfer magnitudes (0.9, 1.4, and 3.6) for comparison. Error bars represent 95% confidence intervals. Note that y-axis ranges differ between panels to facilitate visualization of learning and transfer performance.

The LMM revealed a significant main effect of modality, F(1, 482.95) = 36.23, p < .001, suggesting more accurate performance in the line task (M = 0.88, SD = 0.29) than in the time task (M = 0.80, SD = 0.24). There was also a significant main effect of trial bin, F(4, 478.55) = 2.46, p = .045, suggesting that participants improved in accuracy in both conditions and showed significant learning over trials. There was no significant main effect of feedback condition on learning, F(1, 478.55) = 3.61, p = .061, nor any significant interactions among trial bin, modality, and feedback condition, all ps > .20.

Because the omnibus LMM does not directly characterize the functional form of learning trajectories, we performed additional model-fitting analyses separately for each modality and feedback condition to further examine trajectory shape. Logarithmic, linear, and quadratic models were fit separately for each modality and feedback condition. In the explicit feedback condition, significant logarithmic learning trends were observed across both modalities (time task: adjusted R^2^= .033, p=.014; line task: adjusted R^2^= .022, p=.04), suggesting rapid early improvements for both space and time followed by later stabilization. To further compare learning dynamics across modalities, logarithmic slopes were compared between the time and line tasks. No significant difference in logarithmic slopes was observed, b = 0.008, SE = 0.025, t(271) = 0.31, p = .756, suggesting similar rates of learning across modalities. In the time task with explicit feedback, logarithmic, quadratic, and linear models all significantly fit the data (adjusted R^2^s=.023 ™.033, ps < .05). However, the logarithmic model provided a simpler characterization of the learning trajectory, as reflected by lower AIC and BIC values than the linear and quadratic models (logarithmic: AIC = ™54.0, BIC = ™44.9; linear: AIC = ™52.5, BIC = ™43.4; quadratic: AIC = ™52.7, BIC = ™40.6). In contrast, no significant learning trends were observed in the no feedback condition for either task, as all model fits were nonsignificant (all ps ≥ .265) and explained little variance (all adjusted R² ≤ .005). Full model comparison statistics are reported in the Supplementary Materials (Table S1).

To determine whether time and space productions ultimately reached the target ratio of 1.0 across training, one-sample t-tests were conducted separately for each modality, feedback condition, and trial bin using the Holm correction for multiple comparisons. For the time task, productions in both feedback conditions remained significantly below the target ratio across all trial bins (all Holm-corrected ps < .001), suggesting that time performance failed to fully reach the target regardless of feedback condition, even after 25 training trials. Furthermore, direct comparisons between the feedback and no feedback conditions were not significant after Holm correction (all ps > .07), suggesting that explicit feedback was insufficient to produce reliable improvements in temporal accuracy during training.

In the line task with explicit feedback, only the first trial bin (Trials 1–5) significantly differed from the target ratio, t(29) = ™3.36, p = .002, whereas productions during later trial bins no longer differed significantly from 1.0 (all ps > .083). This suggests successful calibration across training in the line task and near ceiling performance. In contrast, in the line task with no feedback, productions were significantly below the target ratio across all trial bins except the first (Trials 1–5), t(24) = ™1.68, p = .107, where relatively high variability may have contributed to the nonsignificant effect.

### Feedback differentially affects performance accuracy during testing and transfer

Although our analyses of learning blocks suggested no effects of feedback, or at least ambiguous effects of feedback, we next examined performance during the testing phase. Because the testing phase immediately followed training, it provided an opportunity to assess whether any benefits of feedback acquired during learning persisted after feedback was removed. We conducted a separate LMM on the temporal error ratio to examine the effects of modality (time vs. line; within-subjects) and feedback condition (explicit feedback vs. no feedback; between-subjects) on accuracy averaged over blocks (see Fig 2B). This analysis revealed a significant main effect of feedback condition, b = ™0.145, SE = 0.068, t(87.36) = ™2.15, p = .034, such that participants in the explicit feedback condition (M = 0.95, SD = 0.22) showed higher production ratios during the test period, closer to the target value of 1.0, than those in the no feedback condition (M = 0.77, SD = 0.29). There was also a significant main effect of modality, b = ™0.131, SE = 0.051, t(48.15) = ™2.55, p = .014, with participants producing ratios closer to 1.0 in the line task (M = 0.96, SD = 0.25) than in the time task (M = 0.79, SD = 0.26). The interaction between feedback condition and modality, however, was not significant, b = ™0.073, SE = 0.079, t(48.72) = ™0.92, p = .360, suggesting that the effect of feedback on accuracy did not differ across modalities.

The third phase of our experiment involved transfer to new untrained temporal intervals and line lengths. To assess the effects of training and feedback on transfer, we conducted a separate LMM to examine performance on the untrained intervals and distances (0.9, 1.4, and 3.6; within-subjects). We included modality (time vs. line; within-subjects) and feedback condition (explicit feedback vs. no feedback; between-subjects) as additional variables (Fig 2B). The analysis suggested differences in accuracy for different temporal intervals and spatial distances, supported by a main effect of magnitude (F(2, 271.97) = 13.74, p < .001). Follow-up Holm-corrected pairwise comparisons revealed that transfer performance at 3.6 was significantly lower than at both 0.9 (b = 0.156, SE = 0.037, t(272) = 4.21, p < .001) and 1.4 (b = 0.178, SE = 0.037, t(272) = 4.81, p < .001). Performance did not differ between the novel transfer magnitudes of 0.9 and 1.4 (b = −0.022, SE = 0.037, t(272) = −0.60, p = .549). There was also a significant main effect of feedback condition, F(1, 57.78) = 6.74, p = .012, indicating overall differences in transfer performance differed between the explicit feedback and no feedback groups.

Specifically, participants in the feedback condition (M = 1.07, SD = 0.39) showed higher ratios (i.e., greater overproduction) overall compared to the no feedback condition (M = 0.88, SD = 0.38). This pattern was further qualified by a significant interaction between modality and feedback condition, F(1, 279.08) = 7.66, p = .006, indicating that the effect of feedback differed across modalities. Follow-up simple effects analyses revealed that, in the line task, participants in the explicit feedback condition (M = 1.14, SD = 0.42) showed significantly higher ratios (i.e., greater overproduction) than those in the no feedback condition (M = 0.86, SD = 0.39), b = 0.273, SE = 0.078, t(77.6) = 3.51, p < .001. In contrast, no significant effect of feedback was observed in the time task on transfer, with similar ratios in the explicit feedback (M = 1.00, SD = 0.35) and no feedback conditions (M = 0.90, SD = 0.37), b = 0.102, SE = 0.079, t(82.5) = 1.29, p = .202. No other interaction effects were significant (ps > .45).

To examine transfer performance relative to the trained target magnitude (2.8), follow-up comparisons were conducted using the Holm correction for multiple comparisons within each condition (Fig 2B). In the time task with explicit feedback, transfer performance at 0.9 and 1.4 were significantly higher than the trained magnitude of 2.8 (0.9: b = 0.201, t(27) = 3.34, p = .005; 1.4: b = 0.216, t(27) = 5.49, p < .001). The 3.6 interval did not significantly differ from the trained target, b = 0.014, t(27) = 0.52, p = .610. In the time task without feedback, surprisingly, all transfer magnitudes were significantly greater than the trained target condition (0.9: b = 0.323, t(19) = 3.58, p = .002; 1.4: b = 0.299, t(19) = 4.09, p = .002; 3.6: b = 0.103, t(19) = 3.97, p = .002). Similarly, in the line task with explicit feedback, performance ratios at 0.9 and 1.4 were significantly higher than the trained magnitude of 2.8 (0.9: b = 0.192, t(28) = 2.86, p = .016; 1.4: b = 0.233, t(28) = 3.60, p = .004), whereas 3.6 did not significantly differ from the trained target, b = 0.091, t(28) = 1.97, p = .059. These patterns should not necessarily be interpreted as evidence of successful transfer from the trained 2.8 magnitude to other magnitudes. Instead, they may reflect a general tendency for responses to regress toward a central reference value, consistent with central tendency effects that have been widely documented in temporal estimation [35] whereby shorter magnitudes tend to be overproduced and longer magnitudes tend to be underproduced. Although traditionally associated with temporal judgments, a similar pattern was observed in the present spatial task. Full trained-versus-transfer comparisons for all feedback and modality conditions are reported in Table S2.

We further compared the transfer performance with the target ratio 1.0 to see if participants ultimately reached the target using one-sample t-tests with Holm correction applied within each condition. In the time task, the 3.6 transfer magnitude showed a significantly lower ratio than 1.0 under both explicit feedback, b = −0.118, t(29) = −2.58, p = .045, and no feedback, b = −0.236, t(24) = −5.03, p < .001. In contrast, the 0.9 and 1.4 transfer magnitudes did not significantly differ from 1.0 in either feedback condition (all ps ≥ .47). In the line task with explicit feedback, the 1.4 transfer magnitude produced a ratio significantly greater than 1.0, indicating overproduction relative to the target magnitude, b = 0.203, t(30) = 2.62, p = .042. In contrast, the 0.9 and 3.6 transfer magnitudes did not significantly differ from 1.0 (ps > .11). In the line task without feedback, the 3.6 transfer magnitude produced a ratio significantly lower than 1.0, indicating underproduction relative to the target magnitude, b = −0.240, t(27) = −3.50, p = .005, whereas the 0.9 and 1.4 transfer magnitudes did not significantly differ from 1.0 (ps > .13).

Overall, feedback appeared to shift productions of the smaller transfer magnitudes toward higher (more accurate) ratios in both the temporal and spatial tasks. However, the consequences of this shift differed across modalities. In the time task, for which performance at the trained magnitude of 2.8 remained below the target ratio of 1.0, higher ratios tended to bring transfer performance closer to the target. In contrast, because performance at the trained 2.8 magnitude in the line task was already close to 1.0, a similar increase in ratios resulted in overproduction of the smaller transfer magnitudes.

### Feedback promotes early reductions in variability during learning

Training typically works through effects on both accuracy and precision, with changes in precision (variability, measured through the CV) providing important additional insight about how people adjusted their estimates for lines and intervals based on feedback. A separate linear mixed-effects model (LMM) was conducted to examine the effects of trial bin (5 bins; within-subjects), modality (time vs. line; within-subjects), and feedback condition (explicit feedback vs. no feedback; between-subjects) on precision (CV; see Fig 3A). The analysis revealed significant main effects of trial bin, F(4, 485.53) = 20.56, p < .001, and modality, F(1, 487.66) = 61.68, p < .001, indicating that variability changed across training and was overall greater in the time task (M = 0.20, SD = 0.13) than in the line task (M = 0.13, SD = 0.10).

**Fig 3.**
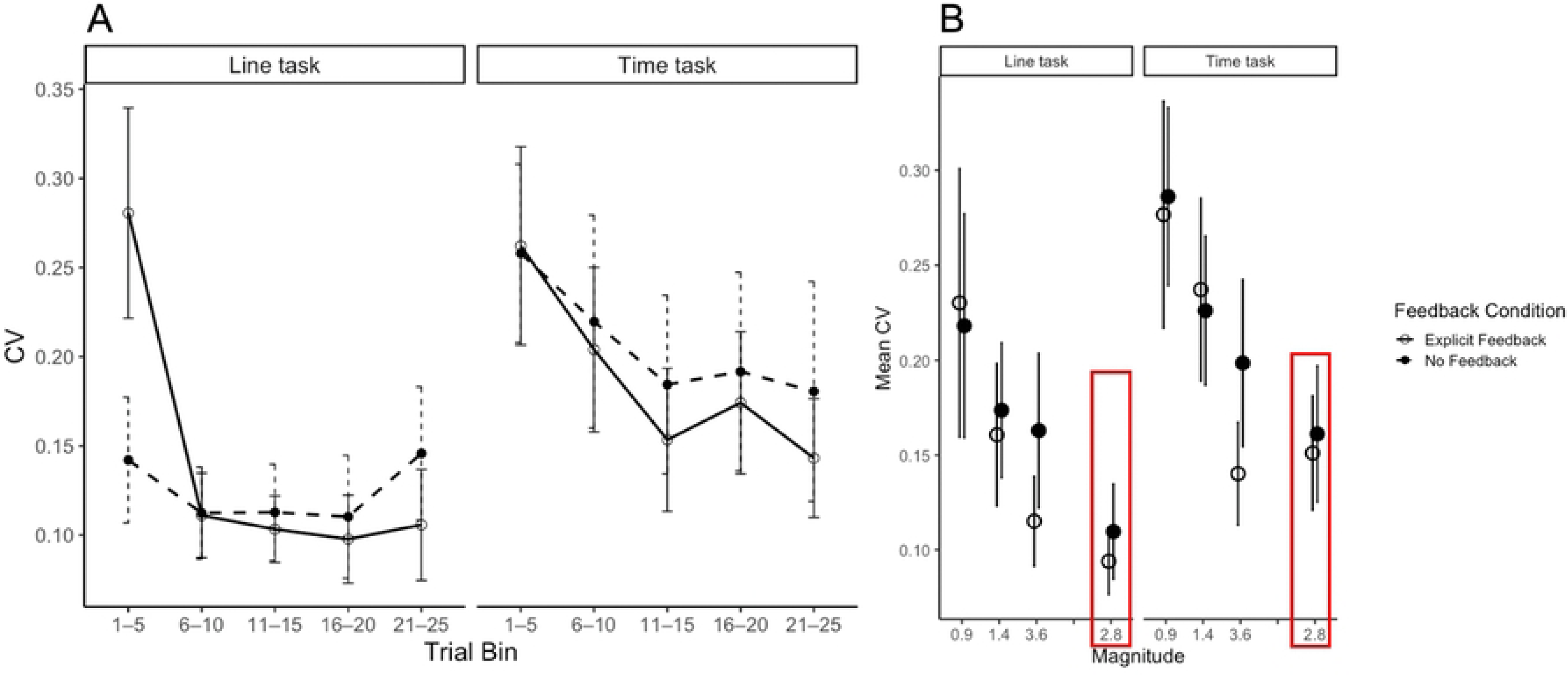
Precision (coefficient of variation; CV) during learning and transfer. (A) Mean CV across training trial bins as a function of modality and feedback condition. The line task is shown on the left and the time task on the right. (B) Mean CV during the transfer phase across magnitudes as a function of modality and feedback condition. Performance at the trained magnitude (2.8) is highlighted by the red box and displayed alongside the untrained transfer magnitudes (0.9, 1.4, and 3.6) for comparison. Error bars represent 95% confidence intervals.

Importantly, a significant interaction between trial bin and feedback condition was observed, F(4, 485.53) = 5.58, p < .001, suggesting that changes in variability across training differed depending on whether explicit feedback was provided in both modalities. Post hoc comparisons revealed that in the explicit feedback condition, CV during the first trial bin (Trials 1–5) was significantly higher than all later trial bins (all Holm-corrected ps < .001), whereas no significant differences were observed among later trial bins themselves (all ps > .32). This pattern suggests that reductions in variability occurred rapidly early in training and subsequently stabilized. In contrast, no significant differences between trial bins were observed in the no feedback condition following correction for multiple comparisons (all Holm-corrected ps > .05).

Similar to the ratio analyses, learning trajectories in CV across trial bins were further characterized separately for each modality and feedback condition using logarithmic, linear, and quadratic models. With explicit feedback, changes in precision in the line task were best characterized by a quadratic model (adjusted R² = .320, p < .001), suggesting a rapid early reduction in variability followed by later stabilization. In contrast, CV in the time task under explicit feedback was best fit by a logarithmic model (adjusted R² = .100, p < .001), indicating more gradual improvements that diminished across training. In the no feedback condition, model fits were substantially weaker across both modalities. The line task showed a weak quadratic pattern (adjusted R² = .022, p = .094), whereas the time task was best characterized by a logarithmic model (adjusted R² = .035, p = .019), indicating a modest but significant reduction in variability. Full model comparison statistics are reported in the Supplementary Materials (Table S3). Together, these findings suggest that feedback produced more structured reductions in variability, particularly in the line task.

Taken together, variability in the spatial task with feedback showed substantial early reductions followed by little additional change, consistent with a quadratic trajectory in CV. In contrast, variability in the temporal task with feedback showed rapid early reductions followed by diminishing improvements across training, consistent with a logarithmic trajectory in CV.

### Feedback has limited effects on precision during testing and transfer

We conducted a separate LMM to examine the effects of modality (time vs. line; within-subjects) and feedback condition (explicit feedback vs. no feedback; between-subjects) on CV during the testing phase (Fig 3B). The analysis revealed a significant main effect of modality, b = 0.057, SE = 0.013, t(42.82) = 4.29, p < .001, with participants showing greater variability in the time task (M = 0.16, SD = 0.078) than in the line task (M = 0.10, SD = 0.050). However, there was no significant main effect of feedback, b = 0.020, SE = 0.019, t(80.64) = 1.04, p = .30, nor a significant interaction between feedback condition and modality, b = ™0.011, SE = 0.021, t(43.93) = ™0.55, p = .59 on CV. These results suggest that participants in the explicit feedback condition (M = 0.12, SD = 0.072) did not exhibit greater precision than those in the no feedback condition (M = 0.14, SD = 0.072).

To further test the effects of training and feedback on transfer precision, another LMM was conducted to examine the effects of transfer magnitudes (0.9, 1.4, and 3.6; within-subjects), modality (time vs. line; within-subjects), and feedback condition (explicit feedback vs. no feedback; between-subjects) on CV during transfer (Fig 3B). The analysis revealed significant main effects of modality, F(1, 285.61) = 23.09, p < .001, indicating that variability was overall greater in the time task (M = 0.23, SD = 0.095) than in the line task (M = 0.18, SD = 0.11). There was also a significant main effect on transfer magnitudes, F(2, 281.10) = 3.94, p < .001, such that CV at 0.9 was significantly greater than at both 1.4, b = 0.053, SE = 0.013, t(281) = 4.11, p < .001, and 3.6, b = 0.099, SE = 0.013, t(281) = 7.59, p < .001. In addition, CV at the 1.4 transfer magnitude was significantly greater than at the 3.6, b = 0.045, SE = 0.013, t(281) = 3.48, p < .001, indicating progressively lower variability with larger transfer magnitudes. No significant main effect of feedback condition was observed, F(1, 57.83) = 0.70, p = .406. Likewise, no significant interactions involving modality or feedback condition were detected, including the interaction between transfer magnitude and feedback condition, F(2, 281.10) = 2.80, p = .062. Together, these findings suggest that variability during transfer differed across modalities and transfer magnitudes but was largely similar across feedback conditions.

To examine transfer variability relative to the trained target magnitude (2.8), follow-up paired comparisons were conducted separately for each modality and feedback condition following Holm correction within each condition (Fig 3B). Across both modalities and feedback conditions, transfer CV at the smaller magnitudes (0.9 and 1.4) was consistently greater than CV at the trained magnitude of 2.8 (all ps <.025), whereas CV at 3.6 did not significantly differ from 2.8 (all ps > .072). Full comparison statistics are reported in Table S4. Together, these results suggest that transfer variability at 3.6 was generally most similar to the trained 2.8 condition across all modalities and feedback conditions.

## Discussion

The present study examined how temporal and spatial performance evolved across an extended period of training, followed by a test and a transfer phase. The study design and analysis put a particular focus on the rate of learning, the effects of feedback, and the generalization of learning to untrained magnitudes. Participants completed temporal interval and line-length production tasks under feedback or no-feedback conditions during the training phase. They were then tested without explicit feedback and were subsequently presented with untrained intervals and line lengths to examine how and whether training on one interval or line segment transferred to new ones. The most salient effects in our results were differences between spatial and temporal performance, although we also found effects on feedback in the test and transfer phases and some modest evidence for transfer to untrained intervals and segments in the transfer phase.

We observed differences in both spatial and temporal accuracy in all phases of the experiment, i.e., training, test, and transfer. Overall, line segment production outperformed temporal interval production during training, an effect which persisted during the test and transfer periods. This advantage for spatial performance manifested early, with spatial learning rapidly approaching the optimal ratio of 1 while temporal learning during training did not achieve this, on average. We also found that precision was consistently higher (lower CV) for spatial compared to temporal production. These findings replicate prior work [32] demonstrating that temporal learning is slower and less accurate than spatial learning but extend these findings to longer training intervals and precision. These findings are consistent with a growing body of literature suggesting that temporal and spatial cognition are not equally learnable or modifiable [21–31], highlighting a fundamental asymmetry between spatial and temporal systems in certain experimental contexts.

The present findings consistently demonstrate that spatial performance is more accurate and precise than temporal performance across all conditions. This distinction may stem from fundamental differences in how temporal and spatial information are represented. Temporal theories such as Scalar Expectancy Theory [15, 16] and the attentional-gate model [36] propose that temporal judgments are constrained by internal sources of noise, including variability in pacemaker processes, attentional fluctuations, and memory distortions. In contrast, spatial information can be externally anchored and directly encoded through sensory input. Because temporal judgments rely more heavily on internal mechanisms such as attention, memory, and internal timing processes [1], they are inherently more susceptible to error and variability. This reliance on internal estimation may help explain why extended training and feedback were less effective at improving temporal performance than spatial performance.

Although temporal and spatial performance differed substantially in their overall levels of accuracy and precision, the model-fitting analyses revealed several notable similarities in how learning unfolded over time. Specifically, both domains exhibited systematic improvements across training and were generally characterized by logarithmic-like learning trajectories in accuracy, with no differences in learning rates. Notably, these logarithmic-like trajectories were characterized by rapid early improvements followed by stabilization, indicating that reinforcement mechanisms may be most influential during the initial stages of practice. A similar pattern was found for precision though spatial precision was better characterized by a quadratic trajectory, suggesting a steeper improvement in consistency than was observed for temporal performance. This pattern is consistent with the rapid adaptation accounts proposed by Simen et al. [10] and Cavdaroglu et al. [11], which suggest that temporal adjustments can occur quickly in response to feedback. Because performance approached an asymptote relatively early in training, however, the present findings provide less support for the Learning-to-Time (LeT) framework, which predicts more gradual improvements through continued reinforcement over extended practice. Importantly, this rapid acquisition was not unique to temporal learning but was also observed during spatial length learning, suggesting that rapid feedback-based calibration may reflect a common characteristic of magnitude learning across domains.

The effects of feedback on learning of temporal intervals and spatial segments were more variable across the different phases of the experiments but also showed some consistent patterns. Although we did not observe differences in accuracy between the feedback and no-feedback conditions during training, a feedback-related advantage emerged during the subsequent testing phase. This was highlighted by a main effect of feedback during the test phase, suggesting that both temporal and spatial production benefited from feedback. Feedback effects may not have emerged during training, likely because participants were actively incorporating this information into their estimates for the single interval and segment distances being acquired. Consistent with this idea, precision did show significant effects of feedback early during training, with participants in the feedback condition exhibiting substantial changes between the initial trial bin and subsequent bins for precision. These findings suggest an active calibration process during training, with the resulting calibration leading to improved accuracy during the subsequent test phase. These findings are consistent with previous work (e.g., [8–9, 37]), which demonstrates that feedback plays a critical role in error correction and the reduction of temporal variability.

The benefits of feedback also persisted into the transfer phase, particularly during the line task. As highlighted by a feedback and modality interaction effect, participants in the feedback condition showed greater accuracy than those in the no-feedback condition during the line task. For the line production task, feedback resulted in response ratios that were closer to the target for the larger transfer magnitude (3.6), whereas the shorter transfer magnitudes (0.9 and 1.4) showed overproduction relative to the target. Across both the line and time tasks, responses to the larger transfer magnitude (3.6) appeared closer to the trained magnitude (2.8) than did the shorter transfer magnitudes. Overall, these findings suggest that feedback-supported learning may transfer to untrained magnitudes, with the clearest benefit in spatial production. In terms of precision, variability generally decreased as magnitude increased, with the 3.6 transfer magnitude exhibiting precision comparable to that observed at the trained 2.8 magnitude. One possible explanation is that this pattern reflects a central tendency effect [35]. Specifically, shorter magnitudes tended to be overproduced, whereas the longer magnitudes tended to be underproduced, reflecting a bias toward a central reference value. Alternatively, because 3.6 was the transfer magnitude closest to the trained 2.8 target, the relatively preserved performance at this magnitude may also reflect some degree of generalization to similar timescales (see review [38]). Central tendency and the closer proximity of the trained vs. transfer magnitude of 3.6, however, can only explain a subset of our findings with regard to transfer and feedback. Importantly, spatial learning appeared to benefit more directly from feedback and did show stronger evidence for transfer, as indicated by a modality by feedback interaction.

The present findings are broadly consistent with those reported by Teng and Isham [32], while extending them in several important ways. Both studies demonstrated that feedback facilitates learning and that temporal and spatial magnitude production differ in their learning outcomes. Yet, because Teng and Isham [32] employed only a brief training phase, it remained unclear whether the observed differences reflected only the early stages of learning. In other words, would temporal learning catch up to spatial learning with extended training? The present study addressed this question by extending the training phase and examining learning trajectories over time. It showed that although temporal and spatial learning followed similar trajectories, characterized by rapid initial improvement followed by asymptotic performance, temporal learning did not ultimately catch up to spatial learning, as it failed to reach the target ratio of 1.0. Furthermore, the present study extends previous work by examining both accuracy and precision throughout learning. Incorporating measures of variability provided a more comprehensive characterization of learning, revealing that spatial production was not only more accurate but also more precise than temporal production, and that feedback affected accuracy and precision differently.

One limitation of the present study is the absence of a pre-training assessment of transfer performance. Because transfer magnitudes were only tested after training, it is difficult to determine the extent to which the observed transfer patterns reflect learning-related generalization versus preexisting characteristics of magnitude estimation, such as the central tendency effect. As a result, the present design cannot fully disentangle the contributions of training and baseline performance to transfer outcomes. Future studies incorporating both pre- and post-training transfer assessments would allow for a more direct evaluation of how learning influences transfer across magnitudes. Nonetheless, the comparison of feedback and no-feedback conditions during the transfer phase suggests that feedback had some benefits to transfer, particularly spatial learning.

Another important consideration is the distinct learning processes observed between the time and line tasks, suggesting that they may rely on different underlying mechanisms. While our experimental design aimed to equate the time and line tasks as closely as possible, this may not have been sufficient to allow for a fully direct comparison. One limitation is the potential difference in participants’ familiarity with spatial estimation, particularly in a visually based paradigm. Spatial tasks may be more intuitive or relatable, potentially giving rise to a performance advantage in magnitude production. Future research could address this by incorporating alternative modalities that favor temporal processing (e.g., auditory-based paradigm) to examine whether parameters such as accuracy, precision, and learning rate related to learning specificity and transfer are affected.

## Conclusion

The current study examined how temporal and spatial learning unfolds over time by comparing temporal and spatial magnitude production under different feedback conditions using both trained and untrained targets. Overall, the findings demonstrate that temporal and spatial learning share some similar learning trajectories but differ substantially in their ultimate levels of accuracy and precision, with spatial performance consistently exhibiting greater accuracy and precision than temporal performance. Feedback facilitated learning by improving precision early during training and accuracy during the subsequent test phase, whereas transfer performance reflected both learning-related generalization and broader biases in magnitude estimation. Together, these results provide insight into the mechanisms underlying temporal and spatial learning and highlight important distinctions in how these domains respond to training and feedback.

## Acknowledgments

We would like to thank Collin Bush for his help with data collection.

## Author Note

We have no known conflict of interest to disclose.

## Funding and Ethics Statement

This research was supported by the National Institutes of Health 2R01NS076856-11. This study was approved by the UA IRB 1801200418.

